# Achieving large dynamic range control of gene expression with a compact RNA transcription-translation regulator

**DOI:** 10.1101/055327

**Authors:** Alexandra M. Westbrook, Julius B. Lucks

## Abstract

RNA transcriptional regulators are emerging as versatile components for genetic circuit construction. However, RNA transcriptional regulators suffer from incomplete repression, making their dynamic range less than that of their protein counterparts. This incomplete repression can cause expression leak, which impedes the construction of larger RNA synthetic regulatory networks. Here we demonstrate how naturally derived antisense RNA-mediated transcriptional regulators can be configured to regulate both transcription and translation in a single compact RNA mechanism that functions in Escherichia coli. Using in vivo gene expression assays, we show that a combination of transcriptional termination and RBS sequestration increases repression from 85% to 98% and activation from 10 fold to over 900 fold in response to cognate antisense RNAs. We also show that orthogonal versions of this mechanism can be created through engineering minimal antisense RNAs. Finally, to demonstrate the utility of this dual control mechanism, we use it to reduce circuit leak in an RNA-only transcriptional cascade that activates gene expression as a function of a small molecule input. We anticipate these regulators will find broad use as synthetic biology moves beyond parts engineering to the design and construction of larger and more sophisticated circuits.

## INTRODUCTION

RNAs are now understood to play broad regulatory roles across the cell (1). As such, synthetic biologists have sought to use these versatile natural systems to create a diverse array of parts that can regulate many aspects of gene expression including transcription (2–4), translation (5, 6), and mRNA degradation (7–9). Antisense-mediated RNA transcriptional regulators are particularly versatile because they regulate RNA synthesis as a function of an RNA input and thus can be used to create RNA-only genetic circuitry (2, 10). RNA genetic circuits have many potential advantages over protein-based circuits including the possibility of leveraging advances in RNA folding algorithms and design rules for part design (11, 12) and their natural fast circuit dynamics (10). However, RNA transcriptional regulators still suffer from low dynamic range in comparison to protein-based regulators. This is a significant barrier to using RNA transcriptional repressors in large genetic circuits because low dynamic range can lead to regulator leak that can propagate through a circuit and break its function. Thus an important challenge for RNA engineering is to improve the dynamic range of RNA regulators so that they can be more effective as elements of synthetic genetic networks.

While there has been great progress in improving the dynamic range of RNA regulators by engineering mechanisms that control a single gene expression process (3, 4, 13), only several studies have explored the idea of engineering multiple genetic control processes for tighter regulation (14–16). Specifically, Morra et al. recently combined transcriptional and translational control with two distinct mechanisms - inducible promoters and orthogonal translational riboswitches - to achieve tight control of fluorescent proteins (14). Horbal and Luzhetskyy also recently used a similar approach to control pamamycin production in *Streptomyces albus* (15). Using RNA engineering strategies, Liu et al. pursued a different approach by combining RNA-mediated translation regulators with leader-peptide transcriptional attenuators to create a hybrid RNA mechanism that uses sequential control of translation then transcription to achieve large dynamic range repression and activation (16). Importantly this study showed that multiple RNA structures can be combined together to regulate multiple aspects of gene expression.

An interesting feature of RNA regulatory mechanisms is that they regulate transcription, translation, and mRNA degradation through the conditional formation of simple hairpin structures at defined positions in mRNAs (17). Specifically, transcriptional terminators repress transcription when they form by causing the polymerase to ratchet off the DNA complex (18, 19), ribosome binding site (RBS)-sequestering hairpins block translation by inhibiting ribosome binding (20, 21), and stability hairpins can block the binding of RNases to control mRNA degradation (22, 23). The common connection between structure and function exhibited by RNA regulatory mechanisms reveals an intriguing possibility of engineering hairpin structures that can regulate multiple control points within a single mechanism.

We sought to use this approach on a mechanism that has already been shown to be useful for engineering a growing number of RNA circuits (2, 10, 24). Specifically, we focused on the pT181 attenuator from the *Staphylococcus aureus* plasmid pT181 (25). In its natural form, the attenuator is an RNA sequence in the 5′ untranslated region of a pT181-encoded mRNA for the plasmid replication protein RepC. Alone, the sense RNA attenuator folds into a structure that allows for transcription of the RepC mRNA. When a cis-encoded antisense RNA, or sRNA repressor, is present, its binding to the sense RNA target causes the sense RNA to fold into a structure that exposes a transcriptional terminator upstream of the RepC coding sequence which represses transcription of the mRNA (Figure 1, Supplementary Figure S1). A number of RNA engineering strategies have utilized the pT181 attenuator as a starting point to create RNA genetic networks and gene expression logics. Earlier studies concluded that the attenuator primarily regulates transcription (26), leading initial engineering efforts to use a transcriptional fusion of the attenuator to create basic RNA transcriptional repressors (2). To construct this transcriptional fusion (26), a portion of the RepC coding sequence was included, followed by a stop codon and a separate ribosome binding site for translation of the downstream gene of interest after the transcriptional decision was made by the attenuator (Supplementary Figure S1). In this configuration, poor repression was observed, which motivated engineering the terminator sequence to increase transcriptional repression from 64% to 85% by the addition of GC pairs (2). This was then used to build a library of orthogonal transcriptional repressors (27), and recently the mechanism was inverted to build RNA transcriptional activators (3). Furthermore, a variety of genetic circuits have been constructed with these orthogonal regulators including logic gates (2, 3), transcriptional cascades (2), and single input modules (10).

Intriguingly, early studies on the natural pT181 attenuator mechanism hypothesized that an AGGAG sequence embedded in the 3’ half of the terminator hairpin was the ribosome binding site for repC (28). This would suggest that the pT181 attenuator could function by occluding the RBS to regulate translation. Later, it was determined that the primary mechanism of repression was transcription by comparing transcriptional versus translational reporter gene fusions (26). However the presence of a near canonical RBS sequence in the 3’ terminator hairpin, spaced 12 nt from the start codon of repC suggests the possibility that the pT181 mechanism may in fact have a more powerful effect on gene expression by simultaneously regulating transcription and translation through the conditional formation of a single compact hairpin in response to interactions with an antisense RNA (Figure 1).

**Figure 1.**
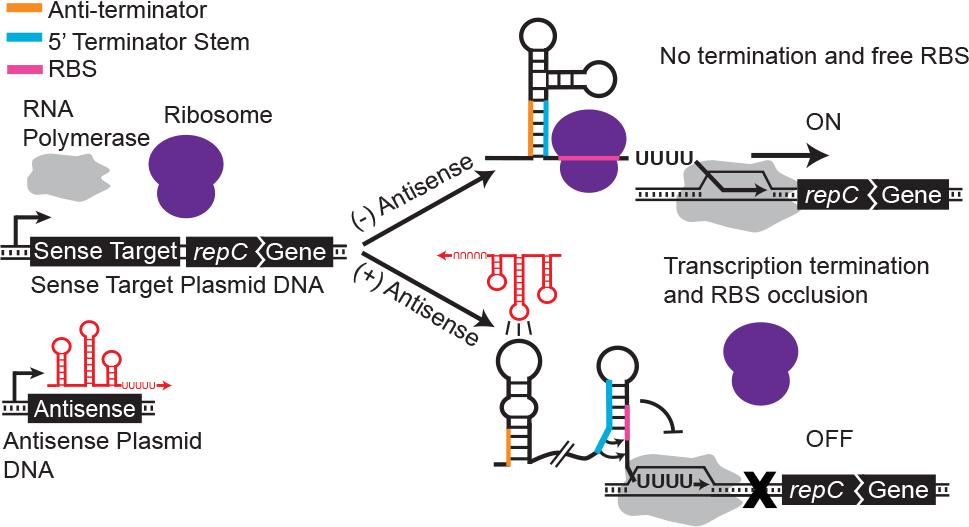
Schematic of the proposed pT181 dual transcription/translation repression mechanism. The pT181 attenuator sense target sequence resides in the 5′ untranslated region and regulates the expression of a downstream gene. The natural attenuator encoded in plasmid pT181 regulates the expression of the *repC* gene (28). Following the attenuator sequence (35), 12 nt of the *repC* gene is included and is translationally fused to the regulated gene of interest. In the absence of antisense RNA (red), the attenuator folds such that the anti-terminator sequence (orange) sequesters the 5′ region of the terminator stem (blue), preventing terminator formation and allowing transcription elongation by RNA polymerase (grey). This structure also contains an exposed ribosome binding site (RBS), which allows ribosomes (purple) to bind and translate the mRNA. Thus in the absence of antisense RNA the attenuator is transcriptionally and translationally ON. When antisense RNA is present, its kissing hairpin interaction with the attenuator sequesters the anti-terminator, thus allowing terminator formation, which prevents downstream transcription. This structure also occludes the RBS inside the 3′ side of the terminator hairpin, which prevents ribosome binding. Thus in the presence of antisense RNA the attenuator is transcriptionally and translationally OFF. A transcriptional regulatory version of the mechanism used in previous engineering (2) is shown in Supplementary Figure S1.

In this work, we show that antisense-mediated repression of gene expression can be improved by utilizing the native RBS and thus the natural dual transcriptional/translational capabilities of the pT181 attenuator. When configured as a translational fusion, we show we can increase the repression of a fluorescent reporter protein from 85% (+/− 3.4%) to 98% (+/− 0.4%) in Escherichia coli. The success of this strategy led us to utilize it to improve the fold activation of a small transcription activating RNA (STAR) system based on the pT181 hairpin from 10 fold (+/− 3.7) to 923 fold (+/− 213). Our next goal was to create a library of orthogonal dual control repressors. To dothis, we applied this strategy to previously published orthogonal pT181 mutants and fusions that functioned at the transcriptional level (27). Interestingly, this library of repressors showed significant cross-talk, indicating that the dual control system breaks orthogonality, likely by increasing theopportunity for non-cognate antisenses to bind and induce translational repression. To mitigate this, we engineered a minimal antisense RNA that greatly improved orthogonality. Finally, to demonstrate that these regulators can be used to fix leak within RNA circuits, we constructed a repressor cascade using the dual control repressor on the bottom level and found that the dual control cascade exhibited reduced circuit leak and a higher dynamic range.

## MATERIAL AND METHODS

### Plasmid construction

Key sequences can be found in Supplementary Table S1. All the plasmids used in this study can be found in Supplementary Table S2 and plasmid diagrams in Supplementary Figure S2. The pT181 repressor and antisense plasmids, the pT181 mutant repressor and antisense plasmids, and the no-antisense control plasmid were constructs pAPA1272, pAPA1256, pAPA1273, pAPA1257, and pAPA1260, respectively, from Lucks et al. (2) The top level of the cascade was the theophylline pT181 mutant antisense plasmid, construct pAPA1306, from Qi et al. (29) The middle level of the cascade was modified from construct pAPA1347 from Lucks et al. (2) using Golden Gate assembly (30). The bottom level of the transcriptional cascade was construct pJBL1855 from Takahashi et al. (10) and the bottom level of the dual control cascade was modified from this construct using Golden Gate assembly. The antisense and repressor plasmids were constructed using inverse PCR (iPCR).

### Strains, growth medium and *In Vivo* end point gene expression

All experiments were performed in *E. coli* strain TG1. Experiments were performed for at least seven biological replicates collected over three separate days. Plasmid combinations were transformed into chemically competent *E. coli* TG1 cells, plated on Difco LB+Agar plates containing 100 µg/mL carbenicillin and 34 µg/mL chloramphenicol and incubated overnight at 37 °C. Plates were taken out of the incubator and left at room temperature for approximately 9 h. Three colonies were picked and used to inoculate 300 µL of LB containing carbenicillin and chloramphenicol at the concentrations above in a 2 mL 96- well block (Costar 3960), and grown approximately 17 h overnight at 37 °C at 1,000 rpm in a Labnet Vortemp 56 benchtop shaker. Six microliters of each overnight culture was then added to separate wells on a new block containing 294 µL (1:50 dilution) of supplemented M9 minimal media (1xM9 minimal salts, 1 mM thiamine hydrochloride, 0.4% glycerol, 0.2% casamino acids, 2 mM MgSO_4_, 0.1 mM CaCl_2_) containing the selective antibiotics and grown for 4 h at the same conditions as the overnight culture. Cultures (6-12 µL) were then transferred into a FACS round-bottom 96 well plate with 244 µL of PBS containing 2mg/mL Kanamycin to stop translation. The plate was then read on a BD LSR II using the high throughput setting with the high throughput sampler (HTS). The samples for Figure 2D were transferred to Falcon 5 ml polystyrene round-bottom tubes and analyzed on a BD Aria Fusion.

### Flow cytometry data collection

Data for the following parameters were collected on the BD LSR II: forward scatter (FSC), side scatter (SSC), and SFGFP (31) fluorescence (488 nm excitation, 515‐545 nm emission). Three to ten uL of each sample was measured in high throughput mode. Each sample was required to have at least 5,000 counts, but most had 10,000 to 50,000. Counts were gated in FSC vs. SSC by choosing a window surrounding the largest cluster of cells. SFGFP fluorescence values were recorded in relative channel number (1‐262,144 corresponding to 18‐bit data) and the geometric mean over the gated data was calculated for each sample. Data for Figure 2C was collected on a BD Aria Fusion for the following parameters: forward scatter (FSC), side scatter (SSC), SFGFP fluorescence (488 nm excitation, 530 nm emission), and mRFP Fluorescence (561 nm excitation, 582 nm emission). SFGFP and mRFP fluorescence values were recorded in relative channel number (1‐262,144 corresponding to 18‐bit data) and the geometric mean over the gated data was calculated for each sample. Compensation was calculated automatically by the BD Aria FACSDiva software using the compensation setup feature.

### Flow cytometry data analysis

Data analysis and FACS calibration was performed according to the supplementary info of Lucks et al. (2) Spherotech 8–Peak Rainbow Calibration Beads (Spherotech cat. no 559123) were used to obtain a calibration curve to convert fluorescence intensity (geometric mean, relative channel number) into Molecules of Equivalent of Fluorescein (MEFL) units for SFGFP fluorescence or Molecules of Equivalent Phycoerythrin (MEPE) for RFP fluorescence. For each experiment, data for a set of control cultures was also collected which consisted of *E. coli* TG1 cells that do not produce SFGFP (transformed with control plasmids JBL001 and JBL002). The mean MEFLor MEPE value of TG1 cells without SFGFP or mRFP expression, respectively was subtracted from each colony's MEFLor MEPE value. Mean MEFLor MEPE values were calculated over replicates and error bars represent the standard deviation. For repressors, the OFF level is the MEFLor MEPE of cells containing the sense plasmid and the antisense plasmid and the ON level is the MEFLor MEPE of cells containing the sense plasmid and a no-antisense control plasmid. The percent repression for each antisense RNA/attenuator plasmid combination was calculated by subtracting the OFF level divided by the ON level from 1 (1‐OFF/ON). For activators the ON level is the MEFLor MEPE of cells containing the sense plasmid and the antisense plasmid and the OFF level is the MEFLor MEPE of cells containing the sense plasmid and a no-antisense control plasmid. The fold activation was calculated by dividing the ON level by the OFF level (ON/OFF).

### *In Vivo* bulk fluorescence time course experiments

Strain, transformation, and media were all the same as for end point experiments described above, except 25µg/mL of kanamycin was used in addition to the other selective antibiotics because the cascade is encoded by three plasmids. Transformation plates containing *E. coli* TG1 cells transformed with three cascade plasmids (Supplementary Table S2) were taken out of the incubator and left at room temperature for approximately 3 h. Three colonies were picked and used to inoculate 300 µL of LB containing selective antibiotics in a 2 mL 96‐ well block (Costar 3960), and grown approximately 17 h overnight at the same conditions as described for an end point experiment. Twenty microliters of each overnight culture was then added to separate wells on a new block containing 980 µL (1:50 dilution) of supplemented M9 minimal media (as mentioned above) containing the selective antibiotics and grown for 4 h at the same conditions as the overnight culture. The optical density (OD, 600 nm) was then measured by transferring 50 µL of culture from the block into a 96‐well plate (Costar 3631) containing 50 µL of phosphate buffered saline (PBS) and measuring using a Biotek Synergy H1m plate reader. The cultures were diluted into 1ml of fresh M9 minimal media to an optical density of 0.015 and grown for four hours. Then theophylline was added to the theophylline condition to a final concentration of 2mM. Every 30 min for the next 4 h, 50 µL from each of the fresh cultures was removed from the 96‐well block and transferred to a 96-well plate (Costar 3631) containing 50 µL of phosphate buffered saline (PBS). SFGFP fluorescence (FL, 485 nm excitation, 520 nm emission) and optical density (OD, 600 nm) were then measured at each time point using a Biotek Synergy H1m plate reader.

### Bulk fluorescence data analysis

On each 96‐well block, there were two sets of controls; a media blank (M9 alone) and *E. coli* TG1 cells that do not produce SFGFP (transformed with control plasmids JBL001, JBL002, and JBL1856). The block contained three replicates of each control. OD and FL values for each colony at each time point were first corrected by subtracting the corresponding values of the media blank at that same time point. The ratio of FLto OD (FL/OD) was then calculated for each well (grown from a single colony), and the mean FL/OD of TG1 cells without SFGFP at the same time point was subtracted from each colony's FL/OD value to correct for cellular autofluorescence (error shown in Supplementary Figure S3). Experiments were performed for nine biological replicates collected over three separate days. Day 1 is shown in Figure 6 while all three days are shown together in Supplementary Figure S3.

## RESULTS

### Regulating both transcription and translation with a single RNA structure improves dynamic range

We first sought to evaluate the performance of the dual control repressor by configuring it as a translational fusion with a downstream reporter gene (Figure 1). Because the terminator hairpin contains a canonical RBS in its 3’ half, we would expect this configuration to regulate both transcription *and* translation of the downstream gene. Specifically, in the presence of antisense RNA, the formation of the terminator hairpin should both repress transcription of the downstream gene, as well as occlude the initiation of translation of any mRNA transcripts that were extended due to imperfect termination efficiency. Thus we expected the dual control translational fusions to exhibit lower OFF levels than the transcription-only regulators.

In previous work, a translational fusion of the pT181 attenuator to the *lacZ* gene exhibited 62% repression in the presence of an antisense RNA as measured by Miller assays (26). Since the terminator of the pT181 system had been previously engineered to increase transcriptional repression, we began by assessing the observed antisense-mediated repression of both the natural and engineered terminator using a translational fusion between *repC* and an SFGFP reporter gene (Figure 2A and Figure 2B). To characterize attenuator function, plasmids were constructed where each attenuator was placed downstream of a constitutive promoter and upstream of the SFGFP coding sequence on a medium copy plasmid. Complementary antisense RNAs were placed on a separate high copy plasmid downstream of the same constitutive promoter (Supplementary Table S1). Each attenuator plasmid was transformed into *E. coli* TG1 cells along with either its cognate antisense or a no-antisense control plasmid (Supplementary Table S2). Individual colonies were picked, grown overnight, subcultured into minimal media and grown until logarithmic growth was reached. Fluorescence was measured for each culture using flow cytometry (see materials and methods). Using this experimental design, we observed a 63% (+/− 7.9%) repression in gene expression for the wild-type transcriptional fusion that increased to 98% (+/− 0.4%) when a translational fusion was used (Figure 2A). A closer examination of the increase in repression revealed that the translational fusion not only decreased the OFF level of gene expression in the presence of antisense, but also increased the ON level in the absence in antisense. We performed the same experiment with the engineered terminator and found an improvement from 85% (+/− 3.4%) repression to 98% (+/− 0.7%) repression (Figure 2B). However, in this case the ON level was reduced for the translational fusion, which could be due to the introduced terminator mutation causing increased spacing between the RBS and the start codon of *repC.* For this reason we chose to continue with the wild type translational fusion repressor.

**Figure 2.**
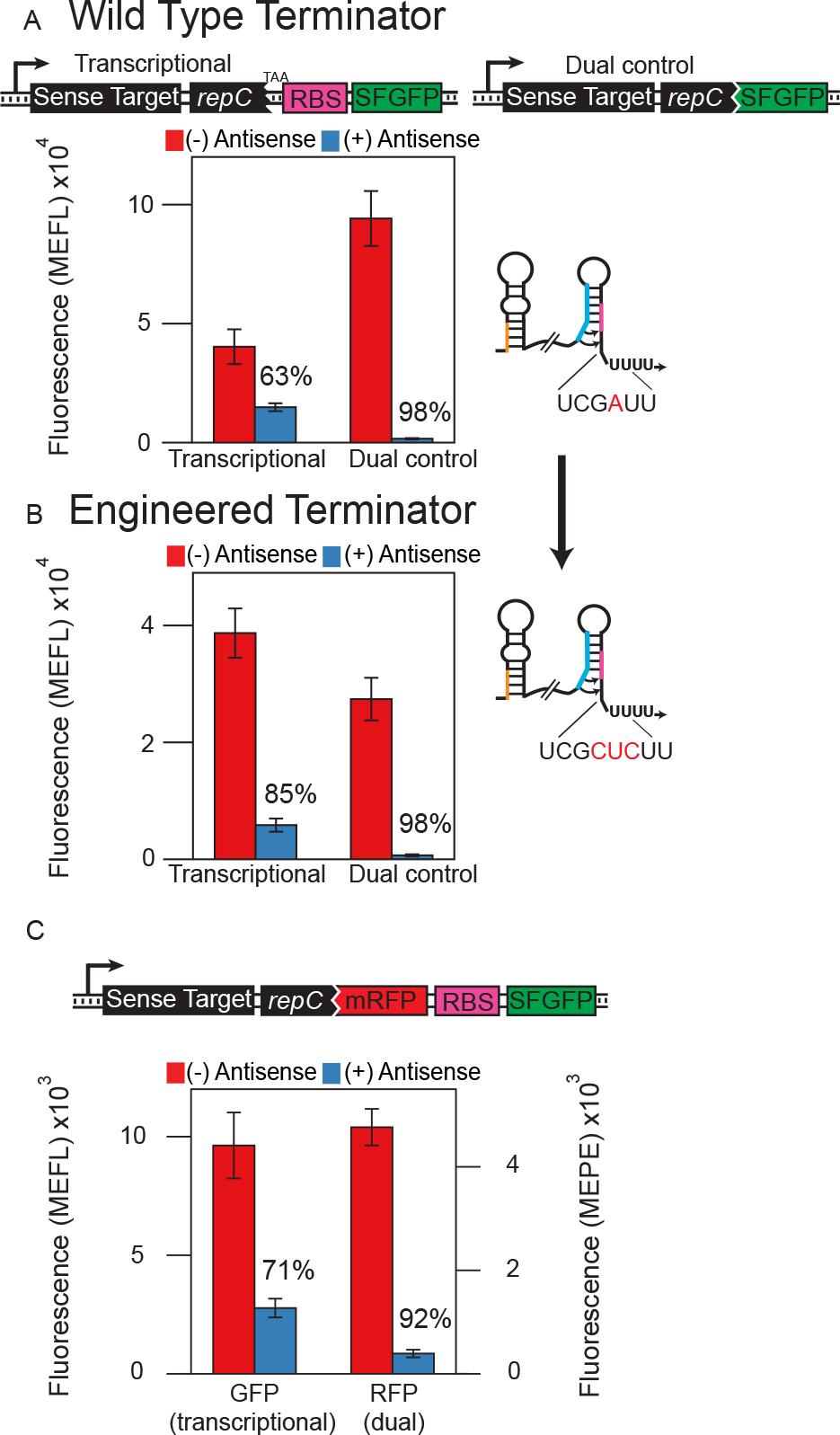
Dual transcription/translation control represses gene expression with higher dynamic range than transcription control in vivo. Functional characterization of the (A) wild type (26), or (B) engineered (2) attenuator configured to repress either transcription (transcriptional fusion) or dual transcription/translation (translational fusion) of an SFGFP coding sequence. Average fluorescence was collected by flow cytometry as Molecules of Equivalent Fluorescein (MEFL) of *E. coli* TG1 cells transformed with a plasmid expressing the indicated attenuator-SFGFP construct and a plasmid expressing the antisense RNA (+, blue) or a control plasmid lacking the antisense sequence (-, red) (Supplementary Table S2). Percent repression is labelled above each construct tested. In both cases the dual control regulator showed 98% repression (50fold), though with a higher ON expression level for the wild type attenuator. Error bars represent standard deviations of at least seven biological replicates. Cartoons highlight differences between the wild type and engineered attenuators, which differ by several bases in the 3′ half of the terminator hairpins. (C) Testing dual control vs. transcriptional control in a two-colour operon construct. The wild type attenuator sequence was translationally fused to an mRFP coding sequence, which was followed by an RBS-SFGFP sequence. In this way mRFP was under dual transcription/translation control while SFGFP was under only transcription control. The construct was tested as in (A) with mRFP fluorescence collected by flow cytometry as Molecules of Equivalent Phycoerythrin (MEPE). RFP was more strongly repressed at 91 % (+/− 1.7%) than GFP at 72% (+/− 5.8%).

We next designed a construct to compare dual transcription/translation control to transcription-only control using a dual reporter protein operon (Figure 2C). In this design, mRFP is translationally fused to the attenuator, while SFGFP is translated from an independent downstream RBS. In this way, we would expect mRFP to be regulated at both the transcriptional and translational levels, while SFGFP would be regulated at just the transcriptional level leading to overall increased repression for mRFP. We transformed cells with the sense target plasmid and the antisense repressor or a blank control plasmid and measured the fluorescence using flow cytometry. As expected, we found that mRFP was repressed more effectively (91% +/− 1.7%) than SFGFP (72% +/− 5.8%). This result also demonstrated that the dual control repressor can be modularly used to regulate different proteins as well as operons.

### The dual control strategy can be extended to a pT181-based activator to dramatically improve fold activation

We next sought to determine if the dual control strategy could be applied to an RNA-based transcriptional activator mechanism derived from the pT181 system. Small transcription activating RNAs (STARs) were recently engineered to activate, rather than repress, transcription in the presence of designed antisense RNAs (3). In the STAR mechanism, the sense target region consists of a transcriptional terminator placed upstream of a target gene which blocks transcription elongation to form the OFF state in the absence of a STAR antisense RNA (Supplementary Figure S4). The addition of a STAR antisense, designed to contain an antiterminator sequence complementary to the 5′ half of the terminator stem, prevents terminator formation, allowing transcription to proceed and gene expression to be ON. Early investigations showed that the pT181 attenuation system could be converted into a STAR by using the terminator sequence from pT181 and an appropriately designed STAR antisense (3). This gave us the opportunity to examine whether a dual control strategy would be effective in the context of gene expression activation.

**Figure 3.**
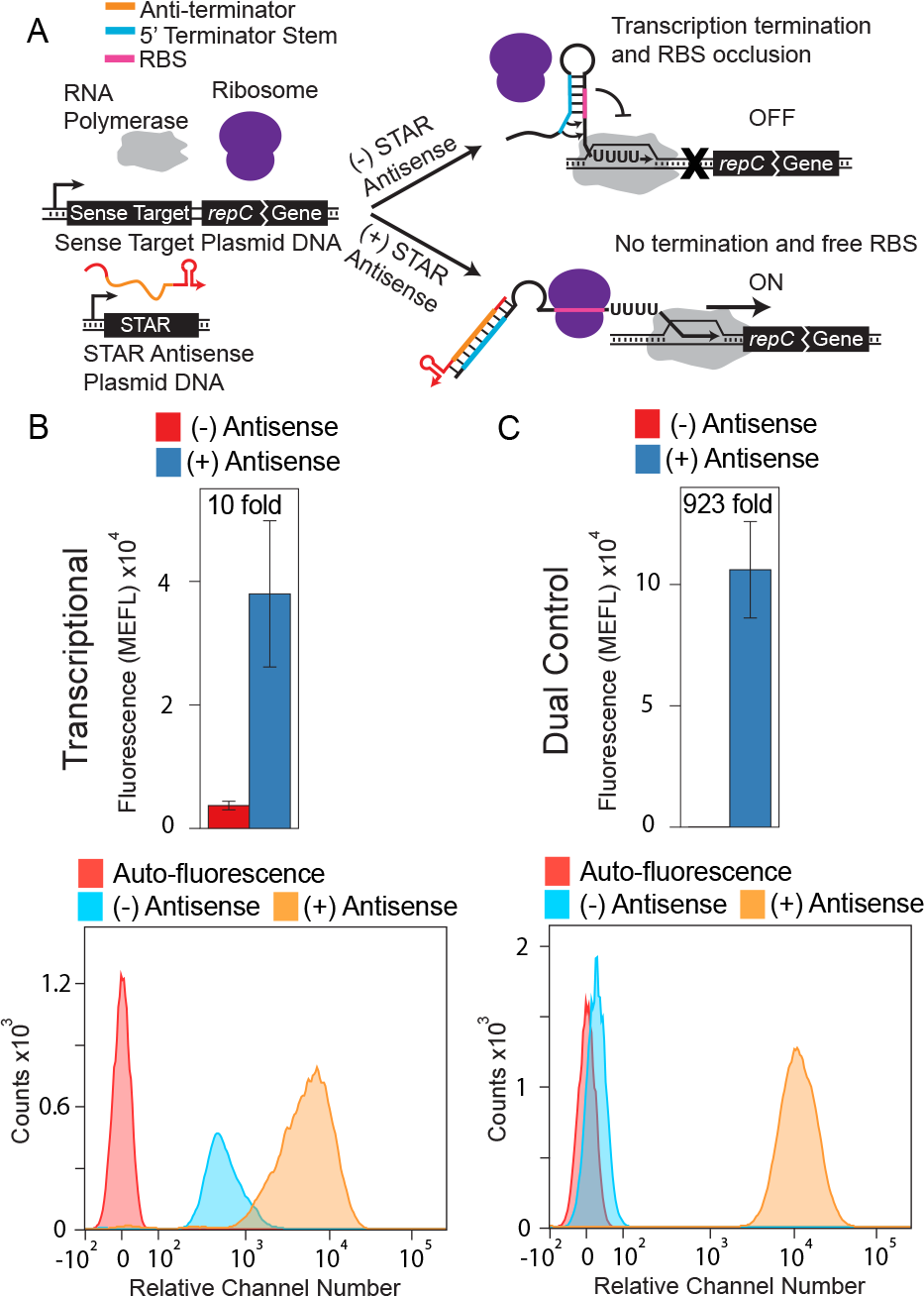
Converting a small transcription activating RNA (STAR) mechanism to a dual transcription/translation activator enhances fold activation. (A) Schematic of the dual transcription/translation activation mechanism. The sense target region consists of the pT181 STAR target region from Chappell et al. (3) followed by 12 nt of the *repC* gene translationally fused to SFGFP. In the absence of the STAR RNA (red/orange), the terminator forms, preventing downstream transcription by RNA polymerase (grey). This structure also occludes the RBS inside the 3′ side of the terminator hairpin, which prevents ribosome binding. Thus in the absence of STAR RNA the mechanism is transcriptionally and translationally OFF. The STAR RNA contains an anti-terminator sequence (orange) complementary to the 5′ half of the terminator (blue). When present, the STAR RNA binds to the terminator, preventing terminator formation and allowing transcription elongation. This structure also exposes the RBS, allowing ribosome binding and translation. Thus in the presence of STAR RNA the mechanism is transcriptionally and translationally ON. The original transcriptional mechanism is shown in Supplementary Figure S4. (B) Functional characterization of a pT 181 STAR that controls transcription. Average fluorescence (MEFL) (top) was collected by flow cytometry of *E. coli* TG1 cells transformed with a plasmid expressing the STAR target transcriptionally fused to an SFGFP coding sequence and a plasmid expressing the STAR RNA (+, blue) or a control plasmid lacking the STAR sequence (-, red) (Supplementary Table S2). Error bars represent standard deviations of at least seven biological replicates. The flow cytometry histogram data (bottom) is plotted on a bi-exponential graph (36). Auto-fluorescence indicates the observed fluorescence distribution from *E. coli* TG1 cells transformed with plasmids lacking activator-SFGFP fusion or antisense (Supplementary Table S2) (C) Functional characterization a pT181 STAR that controls both transcription and translation. Data was collected and plotted as in (B). The dual control strategy increases fold activation from 10 fold (+/− 3.7) to 923 fold (+/− 213) by increasing the ON expression as well as decreasing the OFF expression to near-background auto-fluorescence levels.

To test this, we modified one of the pT181 STARs (Supplementary Table 1) to a translational fusion, making it a dual control activator (Figure 3A). To characterize dual control and transcription-only STAR activator function, each sense target plasmid was transformed into *E. coli* TG1 cells along with either its cognate STAR antisense or a noantisense control plasmid (Supplementary Table S2). Individual colonies were picked, grown overnight, sub-cultured into minimal media and grown until logarithmic growth was reached. Fluorescence was measured for each culture using flow cytometry (see Materials and Methods). The dual control strategy improved transcription-only activation from 10 fold (+/− 3.7) to 923 fold (+/− 213) respectively, due to both a higher ON level and a lower OFF level. Notably the OFF level for the dual-control STAR system was remarkably close to the background cellular autofluorescence level (Figure 3C).

### Multiple dual control regulators can be built using pT181 mutants and chimeras

We next sought to test the orthogonality of the dual control repressors. In addition to having multiple dual control regulators to build genetic circuitry, these regulators must be orthogonal, or only interact with their cognate target. Previous work showed that the original transcription-only chimeric fusions exhibited limited crosstalk between non-cognate antisense/sense target pairs, making them highly We next sought to determine if the dual control strategy could be applied to additional transcriptional attenuators to improve their dynamic range. Multiple orthogonal, or independently acting, pairs of antisense/attenuators are needed in order to build more sophisticated genetic networks. Since a library of orthogonal pT181 transcriptional regulators has previously been engineered (27), we first sought to apply the dual control strategy to these additional regulators. To create orthogonal antisense/attenuator pairs, the library includes several pT181 specificity changing mutants in the first attenuator hairpin that affect antisense recognition, as well as chimeric fusions of the pT181 mechanism with RNA kissing-hairpin interaction regions taken from translational repressors. However, in order to preserve their overall function, the pT181 mutants and fusions share a common expression platform sequence, including the pT181 terminator hairpin, allowing us to make translational fusions to test the dual control strategy in these mutant contexts.

**Figure 4.**
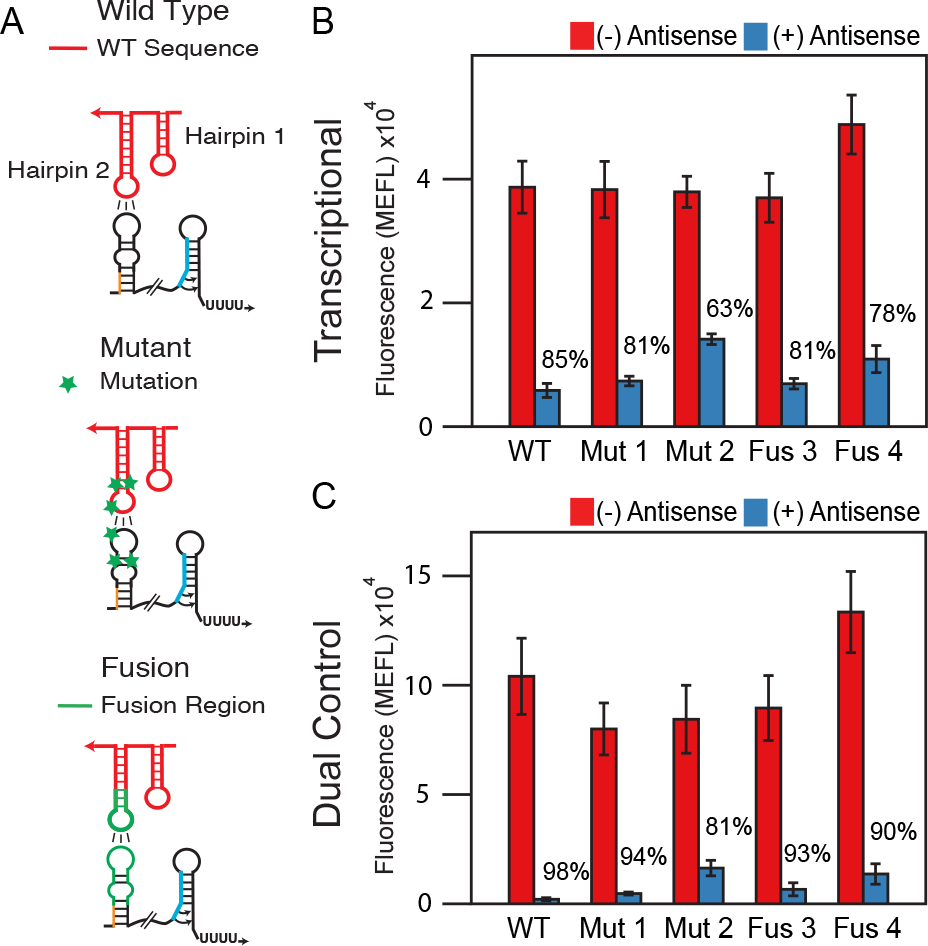
The dual transcription/translation control strategy functions across orthogonal pT181 mutants and chimeras. (A) Schematics of the interactions between the dual control sense target region and the corresponding cognate antisense RNA for wild type, specificity mutants and chimeric fusions engineered to change the specificity of the antisense-attenuator interactions. (B) Functional characterization of the transcriptional wild type pT181 repressor (WT), two mutants (Mut 1,2) (2), and two chimeric fusions (Fus 3,4) (27). Each repressor contained the wild type terminator region depicted in Figure 2A. Functional characterization and data presentation as in Figure 2. Error bars represent standard deviations of at least seven biological replicates. (C) The same as in (B) except with each repressor configured as a dual transcription/translation controller. Using the dual control strategy improves the repression of the transcriptional attenuators.

Additional dual control repressors were characterized as above and compared to the repression observed in the transcription-only regulatory configuration. Specifically, we tested the transcriptional wild type (WT) repressor, the mutant repressors (Mut 1,2) (2), and fusion repressors (Fus 3,4) (27) and found repression percents between 64% and 83% (Figure 4B). We then tested the dual control repressors made from the same attenuators and found that repression increased to between 83% and 98% (Figure 4C) averaging to a 15% increase in repression. As before, these increases in dynamic range come from both a higher ON level and a lower OFF level (Figure 4).

### Orthogonal dual control repressors can be engineered by reducing the antisense RNA sequence

We next sought to test the orthogonality of the dual control repressors. In addition to having multiple dual control regulators to build genetic circuitry, these regulators must be orthogonal, or only interact with their cognate target. Previous work showed that the original transcription-only chimeric fusions exhibited limited crosstalk between non-cognate antisense/sense target pairs, making them highly orthogonal (27). To test this for our dualcontrol repressors, we challenged eachrepressor sense target with all non-cognateantisense RNAs to form an orthogonalitymatrix (Figure 5B, Supplementary FigureS5A). Despite starting from a set of highlyorthogonal transcriptional repressors, weobserved significant crosstalk between thedual control regulators. Earlier work onelucidating the mechanism of antisense-mediated translation repression suggestedthat flanking sequences in the antisenseRNA can form extended interactions with thesense target RNAs (32). We thushypothesized that portions of the antisenseRNAs can be interacting with the sensetarget to repress translation even after thetranscriptional regulatory decision has been made.

**Figure 5.**
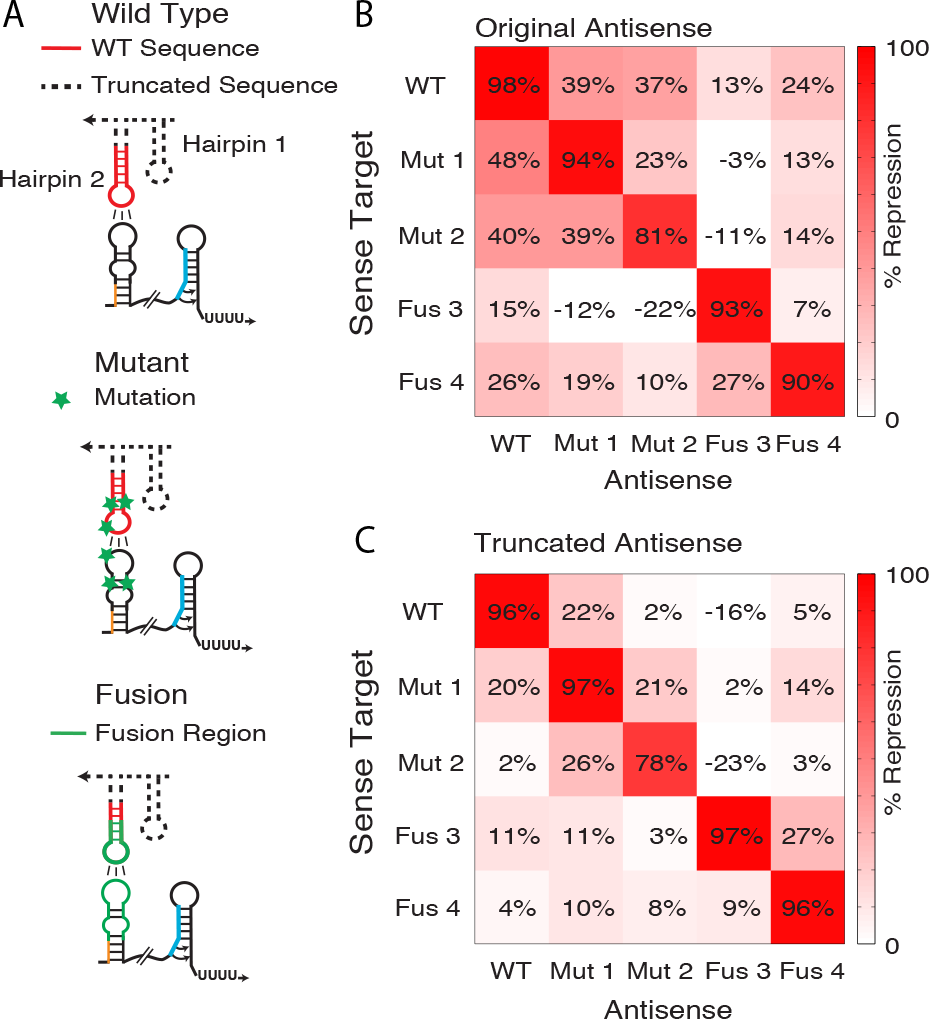
Truncated antisense RNA improves orthogonality between dual transcription/translation RNA repressors. (A) Schematics of the interactions between the dual control sense target region and the corresponding cognate antisense RNA for wild type, specificity mutants and chimeric fusions. Dashed lines show portions of the antisense RNA structure that were truncated to reduce cross talk between pairs of dual transcription/translation control RNA repressors. Hairpin 1 and unnecessary regions (4 nt) at the base of hairpin 2 of the antisense were deleted. (B) An orthgonality matrix showing percent repression observed when sense targets were coexpressed with different full-length antisense RNAs. Each element of the matrix represents the percent repression observed from the indicated antisense/sense target plasmid combination compared to a no-antisense/sense target plasmid condition using functional characterization experiments as in Figure 2. (C) The same as in (B) with truncated forms of the antisense RNAs depicted in (A), showing reduction in repression when non-cognate truncated antisense is present (off diagonal elements). Barplots depicting the data in (B) and (C) are shown in Supplementary Figure S5.

To test this hypothesis, we truncated theantisense RNA sequence to the elementsnecessary for initial RNA-RNA kissing-hairpin interactions that were shown to beessential for the transcriptional regulatorydecision (11). Specifically, hairpin 2 of thepT181 antisense makes contact with the firsthairpin of the sense target region of the attenuator that contains the anti-terminator. We hypothesized that we could remove the antisensehairpin 1 and truncate the stem of hairpin 2 to reduce cross-talk between the dual control repressors(Figure 5A). Using these reduced antisense RNAs, we repeated the orthogonality matrix andobserved that crosstalk was reduced for all non-cognate interactions (Figure 5C, SupplementaryFigure S5B).

### The dual control repressor mitigates circuit leak in an RNA repressor cascade

Finally we sought to test the dual control regulators in an RNA-only circuit context. RNA repressor cascades were the first RNA-only circuits built (2) and have been used to highlight the fast speed of RNA circuitry (10). The repressor cascade also acts as a modular unit that can be built upon to create more sophisticated circuits such as a single input module (SIM) that controls the timing of a sequence of genes in response to a single input (10, 33). However, past attempts at characterizing repressor cascades have revealed circuit leak due to insufficient repression. We therefore sought to fix the leak of an RNA repressor cascade using the dual control repressor. To test this, we built an RNA repressor cascade that activates the expression of SFGFP in response to theophylline (Figure 6A). The cascade consists of three plasmids each expressing one level of the circuit. Without theophylline present, antisense repressor RNA 2 represses sense target RNA 2 and SFGFP expression. When theophylline is added, it activates antisense repressor RNA 1, which is normally non-functional in the absence of theophylline due to a designed interaction between the antisense RNA hairpin and a fused theophylline aptamer (29). In this way, theophylline binding allows antisense repressor RNA 1 to repress antisense repressor RNA 2, allowing SGFP to be expressed. Overall, when theophylline is added to the cell culture, an RNA signal induces SFGFP expression.

**Figure 6.**
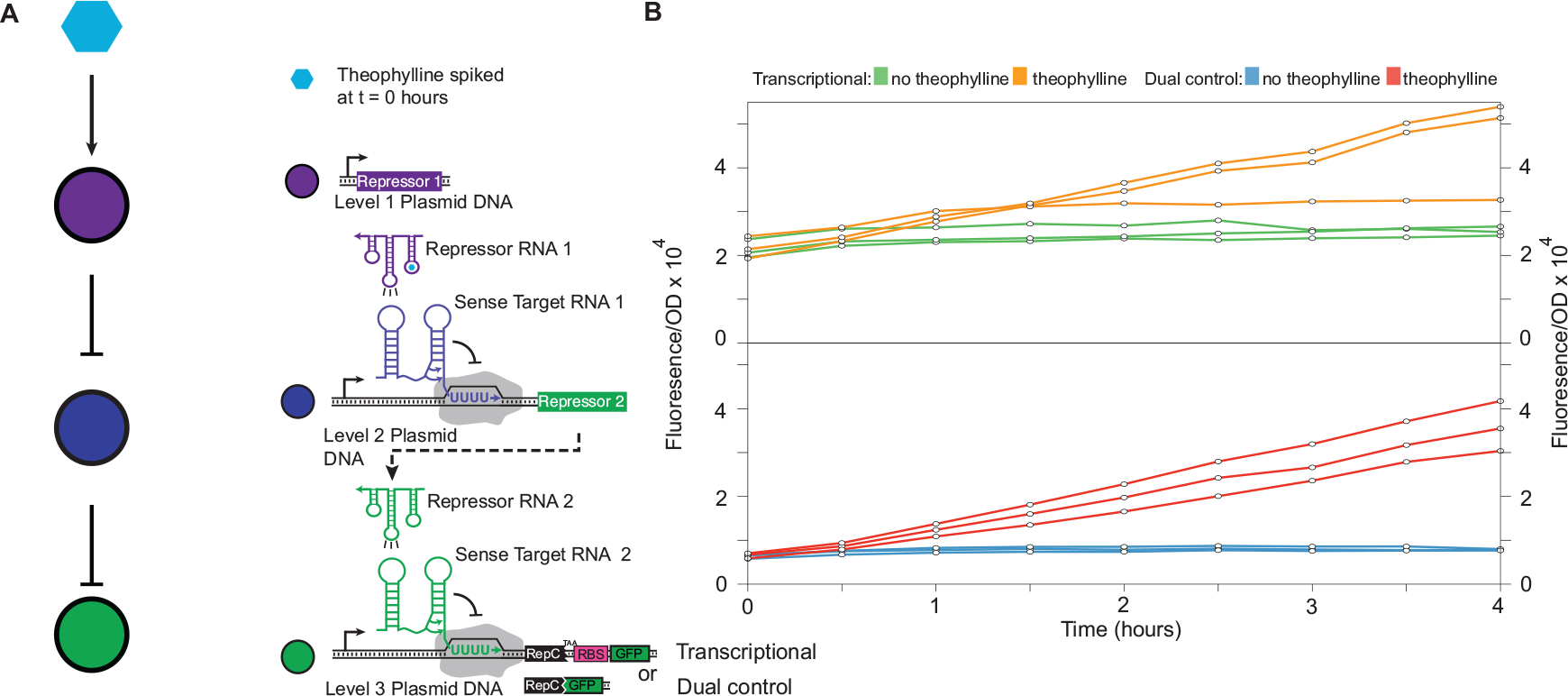
The dual transcription/translation control strategy mitigates leak in an RNA repressor cascade. (A) Schematic of the theophylline activated RNA repressor cascade. The level three SFGFP gene expression is controlled by sense target region 2, which is repressed by repressor RNA 2. Repressor RNA 2 is in turn controlled by the upstream sense target region 1, which is repressed by repressor RNA 1. Repressor RNA 1 is a fusion with a theophylline aptamer (29) that is active only with theophylline bound. Without theophylline, repressor RNA 1 is inactive causing overall repression of SFGFP (OFF). When theophylline is added to the cell culture media, the repressor RNA 1 represses transcription of repressor RNA 2, leading to SFGFP expression (ON). The level three attenuator was configured to regulate SFGFP either transcriptionally, or using the dual transcription/translational control mechanism. (B) Functional time course characterization of the transcriptional and dual control repressor cascades. Three plasmids each encoding one of the circuit levels were co-transformed into *E. coli* TG1 cells, grown overnight and sub-cultured into fresh M9 minimal media for four hours before starting the time-course with a fresh sub-culture (see Methods). After four hours of growth in M9, theophylline (2mM) was added to the media causing SFGFP to be expressed (orange for transcriptional and red for dual control). Time points were sampled every 30 minutes for four hours. Bulk fluorescence and OD600 were measured using a plate reader. The no theophylline condition is shown in green for the transcriptional cascade and blue for dual control. The dual control regulator reduces the overall background fluorescent level while maintaining a similar ON level and thus improves dynamic range. The data shown here are three individual transformants from a single day. Data for three independent experiments performed on separate days are shown together in Supplementary Figure S3. The error associated with subtracting background auto-fluorescence is approximately the size of the points in (B) and is shown in Supplementary Figure S3.

To compare RNA cascades that use either transcription-only or dual control SGFP expression, we performed time course experiments on *E. coli* cultures that contained the cascade plasmids with either the transcriptional or dual control repressor cascade plasmids for the bottom level of the cascade. After incubating overnight in LB media, the cultures were diluted into M9 supplemented media and incubated for four hours. The cultures were then diluted again into fresh M9 media to a consistent OD and incubated for four more hours. From here, we sampled cultures every 30 minutes to measure SFGFP fluorescence and culture OD over time. Theophylline was added to some cell cultures at the beginning of sampling to measure the cascade response (Figure 6B). This experiment was repeated on three separate days, with individual trajectories from the first day shown in Figure 5B and the other two shown in Supplementary Figure S3. As expected, when theophylline was introduced to both the transcriptional and dual control cascades, we observed SFGFP activation that continued throughout the rest of the time course. However, the transcriptional version of the circuit displayed significant leak (Figure 6B, green curves) in comparison to the dual control circuit, which displayed a lower baseline expression (Figure 6B, blue curves) and thus a greater dynamic range. This result demonstrated that dual control repressors can be used in RNA genetic circuits to reduce overall circuit leak.

## DISCUSSION

In this work, we have demonstrated the utility of an RNA structure that regulates both transcription and translation in a single, compact mechanism by showing that it improves dynamic range of antisense RNA-mediated control of gene expression and reduces leak when used in RNA genetic circuits. Specifically, translational fusions between the pT181 attenuator and downstream reporter genes allowed the transcription of these genes to be regulated by the pT181 terminator hairpin and the translation of these genes by the *repC* RBS sequence encoded in the 3’ half of the same hairpin. In this way, the formation of the OFF structure in the presence of a cognate antisense RNA allows gene expression to be repressed at two levels, causing an improvement in repression from 85% (+/− 3.4%) for the transcriptional-only case to 98% (+/− 0.4%) in the dual control case. In addition to decreasing OFF levels in the presence of antisense RNA, this configuration increased the ON level in the absence of antisense RNA. This latter observation could be due to an optimized RNA structural context of the pT181 attenuator that favours efficient ribosome binding. Overall the dual control mechanism significantly improved the dynamic range of RNA regulators over RNA transcriptional repressors and is better than the ~90% repression seen for the best RNA translational repressors (34).

Interestingly, our results are different from those observed by previous studies of the pT181 attenuator (26). Through comparing transcriptional vs. translational fusions of the pT181 attenuator to the LacZ reporter gene, this study observed 62% repression for the translational fusion and 50% for the transcriptional fusion. The lack of significantly different results and the presence of a rho-independent terminator sequence indicated that the attenuator functioned primarily through transcriptional repression. It is possible that our system shows a more significant difference because of the increased sensitivity afforded by our use of SFGFP expression. Nevertheless, our findings strongly suggest that the natural pT181 attenuator system likely regulates at both the transcriptional and translational levels.

In addition to the pT181 dual control repressor, we also engineered a pT181 STAR activator and increased its activation in response to STAR antisense RNA from 10 fold (+/− 3.7) to 923 fold (+/− 213). This improves upon the previously published fold activation of transcriptional STAR regulators (90 fold (3)) and translational toehold regulators (~400 fold (5)). We also showed that this strategy could be expanded to additional pT181 mutants and fusion repressors. Overall this is a significant increase in the number of regulatory tools available for building more sophisticated circuitry with tighter control, which Is particularly useful for situations in which an RNA part with reduced leak is desired.

In order to build robust genetic networks in which the parts act independently and predictably the parts must be orthogonal. However, the initial dual control riboregulators exhibited significant crosstalk. We hypothesized that this was due to additional interactions between the antisense RNAs and the sense target RNAs that caused translation to be repressed even after the transcriptional decision had been made. For a transcriptional decision to be made, the antisense RNAs must interact co-transcriptionally. However, the antisense can still bind the dual control sense target after transcription and affect RBS availability. This would indicate that the modifications between mutants, fusions, and the original pT181 are enough to inhibit crosstalk during transcription but the increased time for antisense-sense target interactions before translation initiation allows shared sequences in non-cognate pairs more opportunity to interact. We therefore decided to reduce the redundant pT181 sequence to reduce the affinity of non-cognate antisense RNAs for sense target regions. By truncating redundant pT181 sequence we greatly improved orthogonality, making it possible to use these dual control repressors in RNA circuitry.

Finally, we used dual control RNA repressors to address a current problem with RNA only circuitry, which is circuit leak that results from parts that do not allow complete repression of their targets. Specifically, we used the dual control repressor in a repressor cascade and found that it reduced circuit leak and background fluorescence.

This work demonstrates a novel RNA that regulates multiple aspects of gene expression in a single compact mechanism, and that displays a dynamic range of gene regulation comparable to protein-based mechanisms. As such, this is another example of how RNAs may be optimized to function as well as proteins. We anticipate as synthetic biology moves beyond the creation of regulator parts libraries and into building more sophisticated networks, RNA regulatory mechanisms such as dual control repressors will find increased use in designing RNA genetic circuits with predictable function.

## ACKNOWLEDGMENTS

We acknowledge Cameron Glasscock (Julius Lucks Laboratory, Cornell University) for his help in reviewing and checking data processing and Eric Strobel (Julius Lucks Laboratory, Cornell University) for insightful discussions of the pT181 mechanism.

